# Pretreatment with yeast derived complex dietary-polysaccharide leads to suppressed gut inflammation, altered microbiota composition and increased immune regulatory short-chain fatty acid production in C57BL/6 mice

**DOI:** 10.1101/719112

**Authors:** Radhika Gudi, Jada Suber, Robert Brown, Benjamin M. Johnson, Chenthamarakshan Vasu

**Affiliations:** Department of Microbiology and Immunology, College of Medicine, Medical University of South Carolina, Charleston, SC-29425

**Keywords:** Yeast β-glucan, gut mucosa, Gut microbiota, short-chain fatty acid, colitis, gut inflammation, immune modulation, immune regulation

## Abstract

**Background:** β-Glucans (BGs), a group of complex non-digestible polysaccharides, are considered to have beneficial health effects due to their immune modulatory properties and are considered as dietary supplements. However, the impact of oral administration of high-pure, well-defined BGs on gut inflammation, and the influence of intestinal microbiota and short-chain fatty acid (SCFA) on the therapeutic effect are largely unknown.

**Objectives:** The aim of this study, using a mouse model of chemical induced colitis, was to investigate the impact of oral administration of high-pure yeast BG (YBG) on the susceptibility to colitis, gut immune function, and structure and function of microbiota.

**Methods:** To determine the impact of oral administration of YBG on colitis susceptibility, eight week old C57BL/6 (B6) mice were pre-treated with YBG (250 μg/mouse/day) and given dextran sulfate sodium (DSS) in drinking water (2.5% w/v) and examined for the symptoms and features of colitis. To assess the effect of oral administration of YBG on gut mucosa and microbiota structure and function, and gut immune regulation, we determined the microbiota composition, fecal SCFA levels, and intestinal T cell phenotype and cytokine secretion. The role of gut microbiota in YBG treatment induced modulation of gut inflammation and immune function were determined in B6 mice treated with broad-spectrum antibiotic cocktail (1 g/L ampicillin, 0.5 g/L vancomycin, 1 g/L neomycin, and 1 g/L metronidazole) in drinking water.

**Results:** Compared to untreated mice, B6 mice that received prolonged pre-treatment with YBG showed diminished severity of different features of DSS-induced colitis including overall loss of body weight (*P*<0.001), shortening of colon (*P*=0.016) and histopathology (*P*=0.01). However, high-pure YBG has no beneficial effect in terms of suppressing colitis severity when consumed only during the disease stage. Compared to untreated controls, YBG pre-treated mice showed higher regulatory T cell (Treg) frequencies (*P*=0.043) in the gut mucosa, a shift in the abundance of gut microbiota towards polysaccharide-fermenting bacterial phyla Bacteroides (*P*=0.049) and Verrucomicrobia (Mean±SD: control=13.0±0.33 vs YBG=10.9.7±0.69) and diminished Firmicutes (*P*<0.001) and Proteobacteria (*P*<0.001), and significantly higher production of SCFA such as acetic acid (*P*=0.016), propionic acid (*P*=0.026) and butyric acid (*P*=0.013). Depletion of gut microbiota in YBG-fed B6 mice using broad spectrum antibiotics caused not only elimination of YBG treatment associated SCFA production and Treg increase, but also profound aggravation of the pathological features of colitis such as loss of body weight (*P*<0.01) and colonic inflammation (*P*=0.04) compared to that of YBG treated control mice.

**Conclusions:** Oral consumption of high-pure BG promotes a healthy gut homeostasis and immune regulation, and minimizes susceptibility to DSS induced colitis in B6 mice in a microbiota (and microbial SCFA) - dependent manner. On the contrary, YBG consumption when gut mucosa and microbiota are compromised not only reverses this protection but also increases the susceptibility to gut inflammation and disease severity, perhaps through its direct interaction with gut immune cells. In conclusion, while YBG consumption may be beneficial for gut health and to prevent gut inflammation in healthy individuals and under intact microbiota, this immune stimulatory dietary supplement may not have any health benefits in individuals with active gut inflammation and could cause adverse effect in those who are on oral antibiotics.

## INTRODUCTION

Dietary approaches have increasingly been considered for modulating immune function, and preventing autoimmunity and other clinical conditions. β-glucans (BGs) are non-digestible complex dietary polysaccharides (CDPs) of varying structures (β-1,3-D-glucan, β-1,3/1,6-D-glucan, β-1,3/1,4-D-glucan and β-1,4-D-glucan) commonly found in plants, cereals, bacteria, yeast, and fungi (1). While BGs can potentiate the function of immune system directly (2, 3), BG-containing preparations have been promoted also as daily dietary supplements to enhance metabolic functions, and as prebiotics (4). On the other hand, clinical studies using YBG preparations reported conflicting outcomes in terms of their potential benefits as dietary supplements (5–9). Studies using animal models to determine the potential of BG preparations and BG containing diets to modulate gut inflammation have also produced conflicting outcomes (10–15). Furthermore, inconsistent results on colitis susceptibility of Dectin-1 (BG interacting receptor) knockout mice have been reported (16, 17).

Differences observed in the impact of BG dietary supplements on host health could be due multiple factors including 1) the purity and/or the structure of the BG employed, 2) degree of direct interactions of those BG preparations with host receptors, 3) possible association between gut microbes and BG induced effects in the host, and differences in the structure and function of basal gut microbiota. Determination of the true effect of oral consumption of BGs on the host, thus, requires systematic pre-clinical studies using well-defined high-pure BGs. Further, studying the direct immune modulatory effect of BGs on gut mucosa and the indirect, microbiota-dependent, effect is necessary to better understand mechanisms of immune modulation induced these dietary complex polysaccharides. Such studies will also help explain the contradictory effects of these agents on the host. For this study, we hypothesized that oral administration of a highly purified BG [β-1,3-linked D-glucose molecules (β-1,3-D-glucan) with β-1,6-linked side chains] from Baker’s yeast (*Saccharomyces cerevisiae*) (YBG) prevents autoimmunity (18, 19) can enhance immune modulation by directly interacting with the gut mucosa and by serving as a fermentation substrate for gut commensals that produce immune regulatory metabolites. Therefore, we examined the impact of oral administration of high pure YBG on gut inflammation employing DSS induced colitis in C57BL/6 (B6) mice as the model, and assessed the contribution of gut microbiota to YBG consumption associated enhanced gut immune regulation and suppression of colitis susceptibility.

## Materials and Methods

### Mice

C57BL/6 (B6) mice were originally purchased from the Jackson laboratory. Foxp3-GFP-knockin (Foxp3-GFP) mice in B6 background were provided by Dr. Vijay Kuchroo (Harvard Medical School, MA). Breeding colonies of these strains were established and maintained in the pathogen-free facility of Medical University of South Carolina (MUSC). Mice were given autoclaved rodent diet (20), which contains 18.6% (wt:wt) protein, 6.2% (wt:wt) fat, and 44.2% (wt:wt) carbohydrates (Harlan Laboratories, Teklad Global 18% Protein Rodent diet no. 2018), and autoclaved water ad libitum. All animal studies were approved by the animal care and use committee at MUSC. Since B6 mice are highly susceptible to DSS induced distal gut inflammation and has been widely used as a model of ulcerative colitis (UC) (21–24), we employed this strain for determining the impact of YBG on DSS induced colitis. While most experiments described in this study used age matched females for control and test groups, our initial experiments showed YBG treatment induced similar protection of male B6 mice from colitis. However, since mortality was as high as 50% by day 10 with control males with DSS colitis, to be able to compare similar number of controls and YBG treated mice side-by-side, we preferred females for this study. B6-Foxp3-GFP report mice that express GFP under Foxp3 promoter and only in regulatory T cell were used in some experiments to visualize and enumerate Tregs by FACS unambiguously.

### YBG and other reagents

Purified YBG (glucan from Baker’s yeast, *S. cerevisiae*), ≥98% pure, was purchased from Sigma-Aldrich and the purification method has been described before (18). This agent was tested for specific activity using thioglycolate-activated macrophages as described before by us and others (25–27). PMA, ionomycin, Brefeldin A, and purified and conjugated antibodies were purchased from Sigma-Aldrich, BD Biosciences, eBioscience, Invitrogen, R&D Systems, and Biolegend Laboratories. In some experiments, intestinal epithelial cells were depleted by Percoll gradient centrifugation (28) or using A33 antibody (Santa Cruz Biotechnology) and anti-rabbit IgG-magnetic beads (29, 30). Oligonucleotide probes for qPCR were custom synthesized by Thermo Fisher. Magnetic bead-based multiplex assay kits were purchased from Invitrogen. Additional details of commercial products used in this study are listed in supplemental table 1.

### YBG treatment

Eight week old B6 mice or B6-Foxp3-GFP mice were given YBG suspension in saline (250 μg/mouse/day) for up to 30 days by oral gavage. Mice that were subjected to colitis induction using DSS, as described below, continued to receive YBG or saline until the mice were euthanized on day 40. In some experiments, YBG and saline treatments were initiated only 1 day before initiating the DSS treatment to determine the effect of treatment initiation at disease stage. This dose of YBG was considered because it is a human relevant dose of BG dietary supplements (500 to 1000 mg/day) equaling approximately 10 mg/kg body weight. Further, as mentioned in our previous report (29), lower doses or short-term treatments do not produce observable immune modulation in the gut mucosa. We also found that even a very short-term (3 day) treatment with higher doses such as 2000 μg/mouse/day produce pro-inflammatory response in the gut mucosa in Dectin-1 dependent manner (29); hence a relatively lower daily dose was considered for this study.

### Induction of DSS colitis and disease evaluation

To induce colitis, mice were maintained on drinking water containing 2.5% (w/v) DSS salt (36,000-50,000 MW) for 5 consecutive days. After 5 days, mice were switched to regular water until euthanasia on day 10, post-DSS treatment initiation. Mice were weighed, and examined for the stool consistency and presence of occult blood every day, and the severities were scored as described before (31). In some experiments, mice were given a cocktail of antibiotics starting 5 days before DSS treatment. Mice were euthanized 10 days post-DSS treatment initiation, intestinal tissues were collected, the colon lengths were measured, and distal colon tissues were subjected to histochemical staining to determine the severity of inflammation. Distal colon tissues were fixed in 10% formaldehyde, 5-µm paraffin sections were cut, and stained with hematoxylin and eosin. Stained sections were analyzed using the 0-6 grading criteria described before (32) for inflammatory cell infiltrate, epithelial changes and mucosal architecture, and the values were averaged to calculate the overall inflammation score of individual mice.

### Immunological assays and qPCR

Cells from the small intestinal lamina propria (SiLP), large intestinal lamina propria (LiLP) and mesenteric lymph node (MLN) were examined for immune cell phenotype and/or cytokine profiles by FACS and multiplex assay respectively as described in our recent reports (29, 30, 33). Briefly: for determining Treg frequencies, freshly isolated cells were stained for CD4 and Foxp3. For determining cytokine positive T cells frequencies, cells were cultured in the presence or PMA, Ionomycin and Brefeldin A for 4 h, before staying for CD4 and intracellular cytokines. For determining secreted cytokine levels, cells were cultured in the presence of anti-CD3 antibody for 24 h and the spent media were subjected to multiplex assay and read using FlexMap 3D instrument (Luminex). SYBR green based qPCR assay was run in a CFX96 Touch real time PCR machine (BioRad) for determining cytokine gene expression levels in the RNA prepared from cryopreserved intestinal tissues.

### 16S rRNA gene targeted sequencing and bacterial community profiling

Total DNA was prepared from the fecal pellets for 16S rRNA gene targeted sequencing and bacterial community profiling as detailed in our previous reports (19, 34, 35) with minor modifications. Briefly, the sequencing reads were fed into the Metagenomics application of BaseSpace (Illumina) for performing taxonomic classification of 16S rRNA targeted amplicon reads using an Illumina-curated version of the GreenGenes taxonomic database which provided raw classification output at multiple taxonomic levels. The sequences were also fed into QIIME open reference operational taxonomic units (OTU) picking pipeline (36) using pre-processing application of BaseSpace. The OTUs were compiled to different taxonomical levels based upon the percentage identity to GreenGenes reference sequences (i.e. >97% identity) and the percentage values of sequences within each sample that map to respective taxonomical levels were calculated. The OTUs were also normalized and used for metagenomes prediction of Kyoto Encyclopedia of Genes and Genomes (KEGG) orthologs employing PICRUSt as described before (37–40). The predictions were summarized to multiple levels and functional categories were compared among control and YBG fed groups using the statistical Analysis of Metagenomic Profile Package (STAMP) as described before (38, 41). STAMP and web-based MicrobiomeAnalyst (42) applications were also employed for visualization and/or statistical analysis of microbiome data.

### Depletion of gut microbiota

Mice were given a broad-spectrum antibiotic cocktail (ampicillin (1 g/l), vancomycin (0.5 g/l), neomycin (1 g/l), and metronidazole (1 g/l) -containing drinking water during the last 5 days of YBG treatment to deplete gut microbiota as described in our recent report (29). Fecal pellet suspensions were cultured under aerobic and anaerobic conditions to assess microbiota depletion (29). In some experiments, antibiotic treatment was continued during colitis induction using DSS.

### Determination of fecal SCFA levels

Fresh fecal samples were collected from individual mice, stored at -80°C, and shipped on dry ice to Microbiome Insights (Vancouver, Canada) for SCFA analysis. SCFA were detected by this service provider using gas chromatography (Thermo Trace 1310), coupled to a flame ionization detector, using Thermo TG-WAXMS A GC Column by following a previously described method (43).

### Statistical analysis

Mean, SD, and statistical significance (*P-value*) were calculated, graphic visualizations were made using Excel (Microsoft), Prism (GraphPad), Morpheus (Broad Institute), STAMP (38, 41) and/or MicrobiomeAnalyst (42) applications. Briefly, unpaired, two-tailed *t*-test or two-tailed non-parametric Mann-Whitney test was employed for most assays where test (YBG fed or microbiota depleted) group samples were compared to that of respective control (saline fed or those with intact microbiota) group, unless specified here. For DSS colitis associated weight loss, stool consistency and stool levels of blood, test group mean of each time point was compared to the respective time-point control group mean by unpaired, two-tailed *t*-test. To compare the number of mice with grade ≤3 and ≥4 colon histopathology scores in test and control groups, two-tailed Fishers’ exact contingency test was employed. Log-rank test was employed for comparing the survival rates of YBG treated mice with that of controls. Statistical analysis of microbiota at genus level as well as the number of sequences representing (predicted) specific metabolic pathways were done in STAMP employing two-sided Welch’s *t*-test and FDR corrected using Benjamini and Hochberg approach, and the heatmap visualization was done in Morpheus. MicrobiomeAnalyst was employed for generating principal component (PC) analysis representing β-diversity (Bray Curtis distance) and α-diversity / species richness / Chao1 estimator plots, and for statistical significances by permutational multivariate analysis of variance and Mann-Whitney approaches respectively. A *p*-value of ≤0.05 was considered significant.

## Results

### Prolonged pretreatment with YBG diminishes susceptibility to DSS induced colitis in B6 mice

Eight week old female B6 mice were orally administered with YBG and subjected to colitis induction using DSS as shown in **Fig. 1A**, and examined for gut inflammation associated features to determine the impact of pre-treatment with YBG on colitis susceptibility. As observed in **Fig. 1B**, YBG treated mice were relatively less susceptible to DSS colitis associated weight loss compared to control mice. Significant difference in the body weight between control and YBG treated groups was observed starting day 6 (*P*=0.027) and peaked at day 10 (*P*<0.001). As shown in **Fig. 1C**, YBG treated mice also showed significant resistance to colitis associated shortening of the colon compared to that of control mice (*P*=0.016). Overall DSS induced histological injury scores of YBG treated mice was significantly lower (*P*=0.016) as compared to that of control group (**Fig. 1D**). The colon of DSS-treated control mice exhibited more severe inflammatory infiltrates and ulceration compared to YBG treated mice. Further, compared to DSS treated control mice, better stool consistency (**supplemental Fig. 1A**) and lesser fecal blood (**supplemental Fig. 1B**) were detected in YBG-fed mice that were treated with DSS at the time-point in which these features were most severe. Of note, prolonged pre-treatment of male B6 mice with YBG produced similar protection to that of B6 female counterparts, but control group showed 50% mortality prior to termination of the experiment (not shown). Further, we also observed that mice that were given YBG only during colitis induction did not show any difference in the severity of colitis associated parameters compared to control mice (**Supplemental Fig. 2**).

**FIGURE 1:**
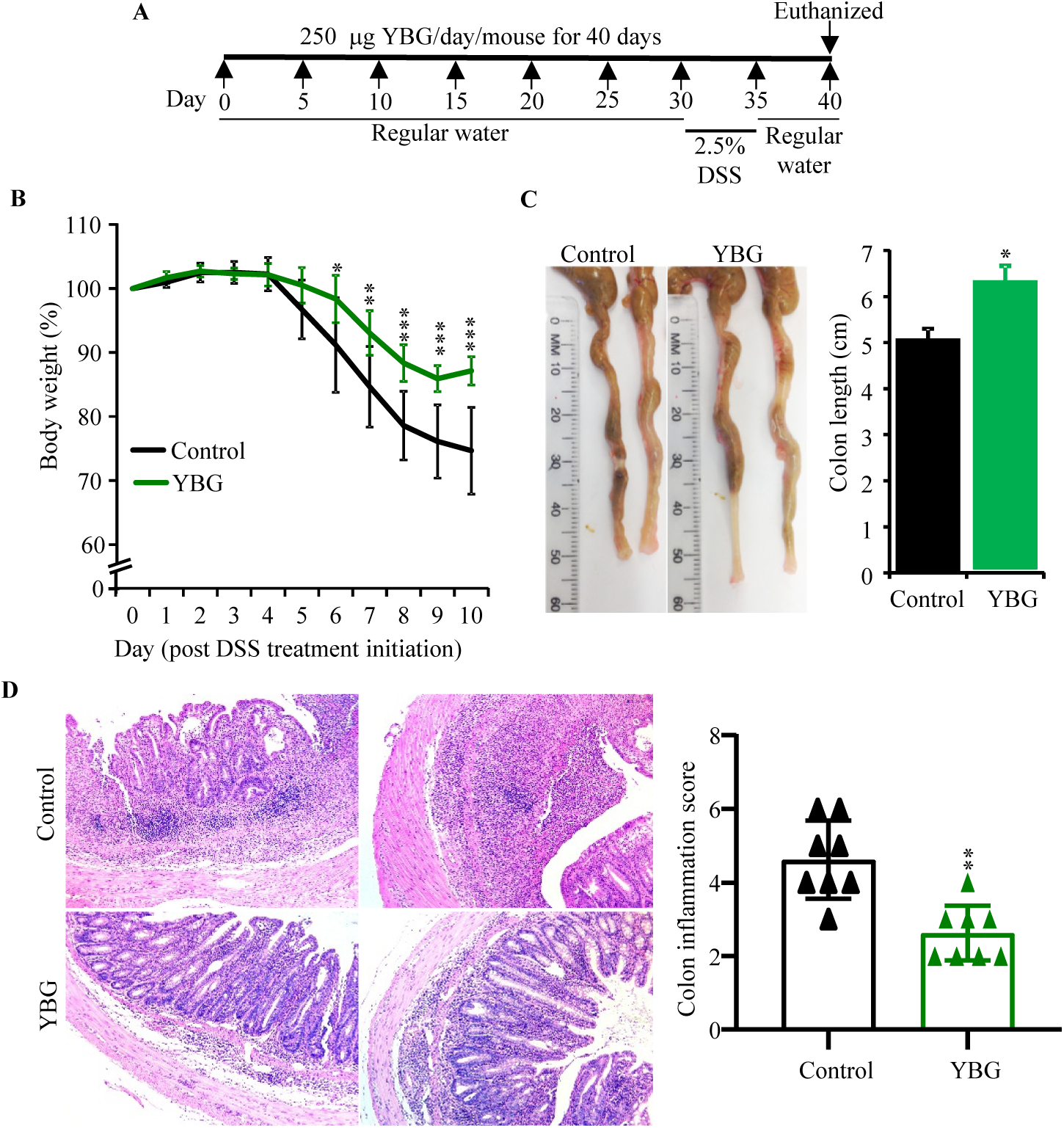
Effect of pre-treatment with YBG (A) on induced colitis associated weight loss (B), colon shortening (C) and colon inflammation (D) in B6 mice. (A) Cartoon depicting experimental design is shown. (B) Body weights of individual animals were measured every day starting at day 30 and changes in percentage of body weights, relative to initial body weight, are shown. (C) Images of representative colons (left panel) and colon lengths (right panel) of euthanized mice are shown. (D) H&E stained distal colon sections were evaluated for inflammation and images of representative sections (left panel) and inflammation severity scores (right panel) are shown. (B, right panels of C, D) Mean±SDs (*n*=8). *, **,*** Different from control, *P*<0.05, *P*<0.01, *P*<0.001 respectively.

### YBG treated and control B6 mice with colitis showed distinct immune characteristics

Since B6 mice that were pretreated with YBG showed less severe colon inflammation, T cells from their gut associated mesenteric lymph nodes (MLNs) were examined for phenotypic properties. As shown in **Fig. 2A** and **supplemental Fig. 3**, Foxp3+ T regulatory cell (Treg) frequencies were significantly higher in the MLN of YBG-fed mice (*P*=0.0043) compared to control mice. FACS analyses revealed significantly higher frequencies of IL10+ (*P*=0.0043), IL4+ (*P*=0.041) and IL9+ (*P*=0.015) and lower IFNγ+ (*P*=0.0087), IL17+ (*P*=0.0022) and IL22+ (*P*=0.0152) T cell frequencies in YBG-fed mice compared to controls. To validate these observations, the levels of cytokines secreted by MLN cell were determined. Multiplex cytokine assay revealed significantly higher secretion of IL10+ (*P*=0.0087), IL4+ (*P*=0.0152) and IL9+ (*P*=0.041) and lower IFNγ+ (*P*=0.0087), IL17+ (*P*=0.041) and IL22+ (*P*=0.026) by MLN cells from YBG fed mice, compared to that of control mice, upon *ex vivo* activation using anti-CD3 antibody (**Fig. 2B**).

**FIGURE 2:**
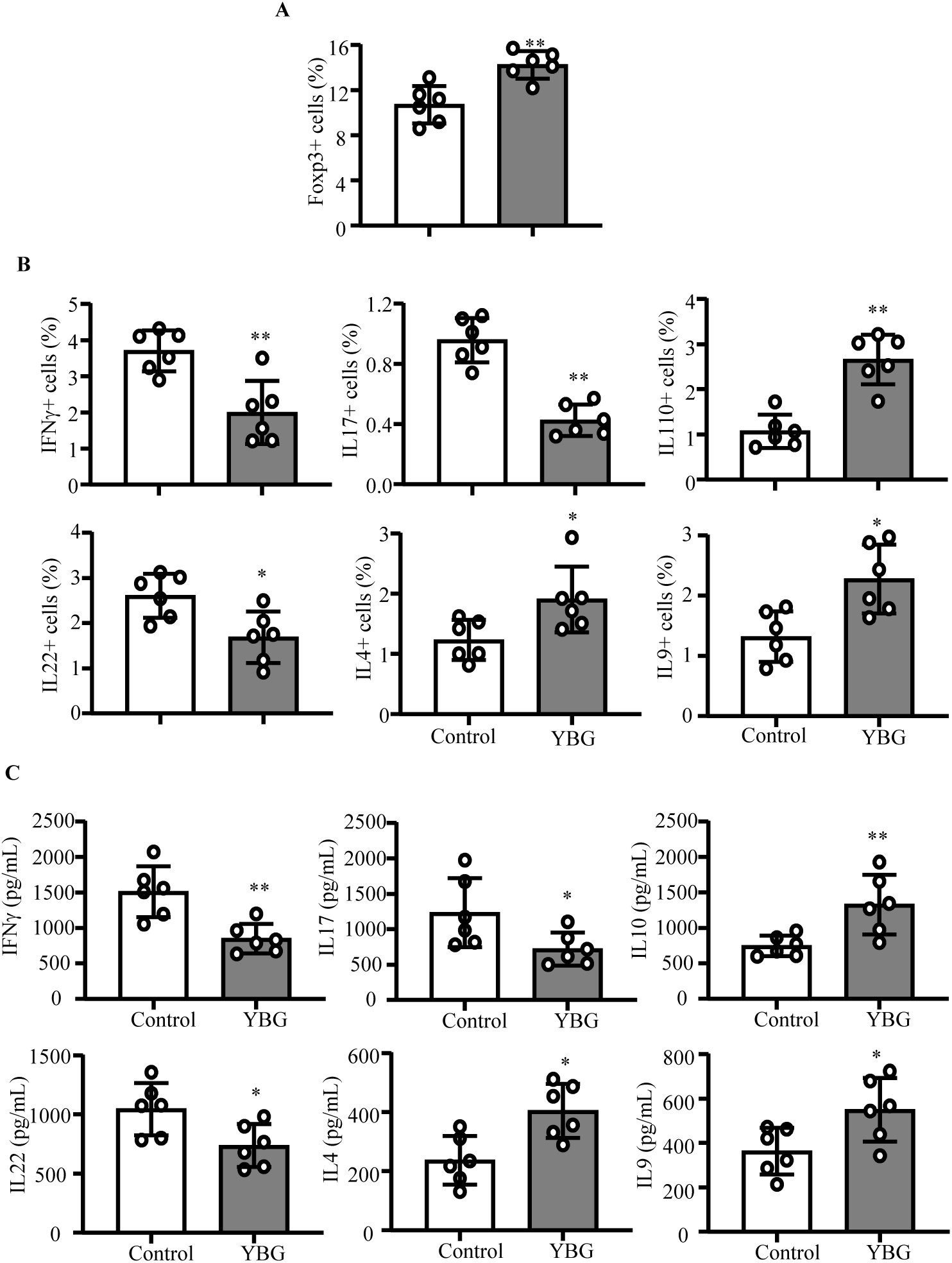
Effect of YBG pre-treatment on Treg (A) and cytokine expressing (B) CD4+ T frequencies in, and cytokine secretion (C) by, MLN cells of B6 mice with DSS induced colitis. (A,B) Frequencies of CD4+ cells in the MLN single cell suspension that express indicated factor determined by FACS are shown. C) Concentrations of indicated cytokines secreted by MLN cells upon *ex vivo* stimulation using anti-CD3 antibody are shown. Mean±SDs (*n*=5) are shown for all panels. *,** Different from control, *P*<0.05, *P*<0.01 respectively.

### Prolonged YBG treatment results in changes in the fecal microbiota composition in B6 mice

Since only prolonged with YBG prior to DSS treatment reduced the susceptibility to colitis in B6 mice, we determined if YBG treatment alters gut microbiota structure and function in them. Compilation of OTU data generated from 16S rRNA sequences of fecal samples to different taxonomical levels showed that, compared to the pre-treatment time-point, abundances of major phyla changed significantly in YBG-fed mice, but not in control mice (supplemental Fig. 4A). YBG treatment resulted in increase in the abundances of Bacteroidetes (*P*=0.049), and Verrucomicrobia (Mean±SD: control=13.0±0.33 vs YBG=10.9.7±0.69) phyla members and decrease in Firmicutes (*P*<0.001) and Proteobacteria (*P*<0.001) in their fecal samples. Interestingly, at genus level, abundances of many major microbial communities are altered significantly after YBG treatment (**Fig. 3A**). Compared to pre-treatment time-point, significant reductions in the abundance of microbial communities belonging to multiple genera including Ruminococcus (phylum: Firmicutes; *P*=0.003), *Lactobacillus* (phylum: Firmicutes; *P*=0.02)*, Oscillospira* (phylum: Firmicutes; *P*<0.001), *Dehalobacterium* (phylum: Firmicutes; *P*<0.006), and *Odoribacter* (phylum: Firmicutes; *P*<0.001) were observed after YBG treatment. On the other hand, abundance of microbial communities belonging to many genera including *Parabacteroidetes* (phylum: Bacteroidetes; *P*<0.005), *Prevotella* (phylum: Bacteroidetes; *P*<0.002), *Sutterella* (phylum: Proteobacteria; *P*<0.007), and *Bifidobacterium* (phylum: Actinobacteria; *P*=0.048) were increased after YBG treatment. Interestingly fecal samples collected after YBG treatment showed lower gut microbial α-diversity / species richness (*P*<0.001) compared to pretreatment time-point (Supplemental Fig. 4B). Further, β-diversity (Bray Curtis distance) analysis of fecal microbiota showed significant distance (*P*=0.007) in the clustering / separation of samples collected before and after YBG treatment (Supplemental Fig. 4C). As shown in **Fig. 3C**, OTU-based prediction of metabolic functions of gut microbes revealed overrepresentation of many pathways including that of carbohydrate metabolism (*P=*0.031), glycan biosynthesis and metabolism (*P=*0.027) and biosynthesis of secondary metabolites (*P=*0.01) in B6 mouse gut microbiota after YBG treatment compared to pre-treatment time-point.

**FIGURE 3:**
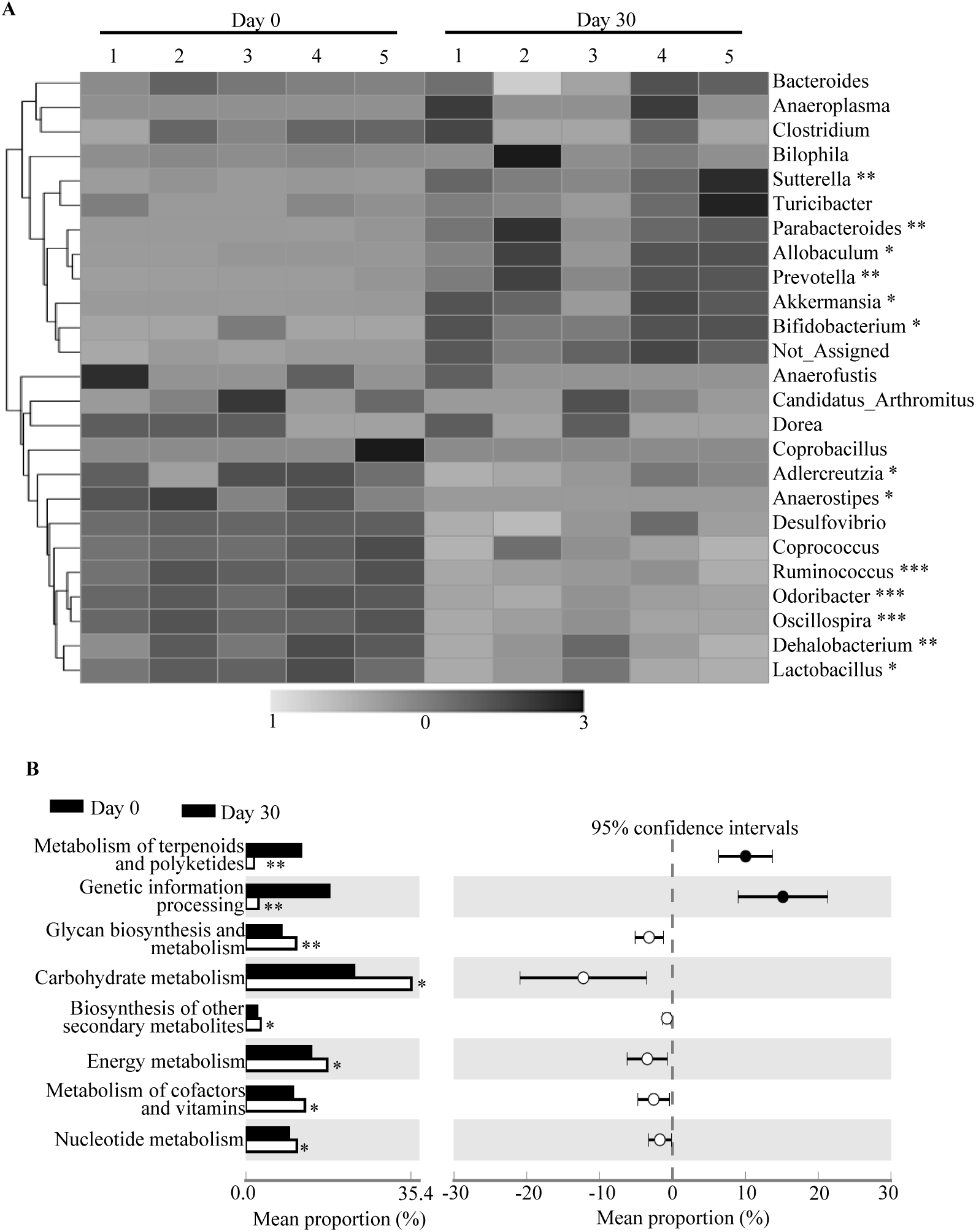
Impact of oral administration of YBG on the composition of gut microbiota at genus level (A) and predictive functions (B) in B6 mice. (A) OTU data of 16S rRNA gene sequences of fecal pellet collected before (day 0) and 30 days after YBG treatment (day 30) were compiled to genus level based upon the percentage identity to reference sequences (i.e. >97% identity) and heatmap was generated using sequence counts of major genera (those with >1 % of total sequences) that showed statistically significant differences. (B) OUT-biom data was used for predicting gene functional categories using PICRUSt application and the level 2 categories of KEGG pathway that showed statistically significant differences are shown. *n*=5 for both panels. *, **,*** Different from control, *P*<0.05, *P*<0.01, *P*<0.001 respectively.

### YBG treatment results in modulation of small and large intestinal immune phenotype in B6 mice

Since YBG treatment altered gut microbiota and suppressed colitis susceptibility in B6 mice, we determined if YBG treatment alone has an impact on gut immune phenotype. Supplemental fig. 5 shows that the primarily small intestine (ileum), but not large intestine (colon), of mice that received YBG for 30 consecutive days expressed significantly higher levels of pro-inflammatory cytokine TNFα (*P=*0.0039) and immune regulatory enzyme Raldh1A2 (*P=*0.001) compared to that of untreated counterparts. On the other hand, significantly higher levels of IL10 expression was detected in both ileum (*P=*0.0017) and colon (*P=*0.0042) of YBG treated mice compared to controls. We then determined the T phenotype of gut mucosa in mice that received YBG for 30 consecutive days by assessing the Foxp3+ T cell frequencies and cytokine production. As shown in supplemental Fig. 6A and **Fig. 4A**, significantly higher frequencies of Foxp3+ T cells were detected in mesenteric LNs (MLNs) (*P=*0.016), small intestine lamina propria (SiLP) (*P=*0.0079) and large intestinal lamina propria (LiLP) (*P=*0.0159) of YBG treated mice compared to controls. Further, compared to that of controls, small intestinal cells from YBG treated mice produced significantly higher amounts of IL10 (*P=*0.016) and IL17 (*P=*0.0044). On the other hand, colonic immune cells of YBG treated mice produced higher amounts of IL10 (*P=*0.0079), but not IL17 or IFNγ.

**FIGURE 4:**
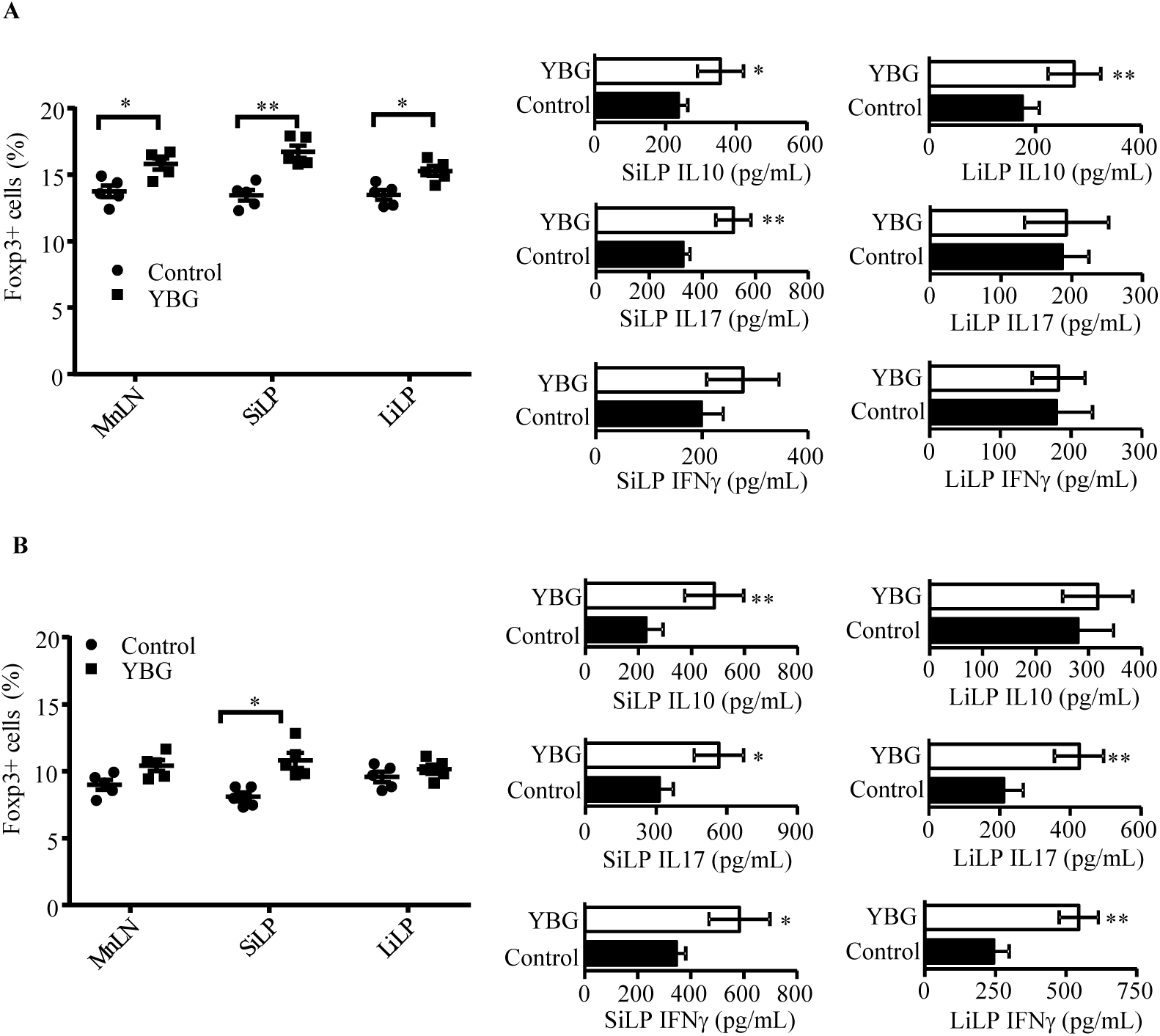
Impact of oral administration of YBG on the Treg frequency and cytokine secretion profile of immune cells in B6 mice with intact (A) and depleted (B) gut microbiota. (A, B) Foxp3+ CD4 cell frequencies in gut associated lymphoid tissues (MLN, SiLP and LiLP) determined by FACS (left panels) and the levels of indicated cytokines secreted by MLN upon stimulation using anti-CD3 antibody determined by multiplex assay (right panels) are shown. Mean±SDs (*n*=5) are shown for all panels. *,** Different from control, *P*<0.05, *P*<0.01 respectively.

We then determined the influence of microbiota on YBG consumption-associated immune modulation in B6 mice after depleting gut bacteria using broad-spectrum antibiotics (Supplemental fig. 7). Depletion of gut bacteria alone caused reduction in Foxp3+ T cell frequencies in both control and YBG recipient mice (supplemental Fig. 6B and **Fig. 4B**), relative to their counterparts with intact microbiota (shown in Fig. 4A). Further, SiLP cells, but not LiLP cells, from microbiota-depleted YBG recipient mice showed relatively higher frequencies of Foxp3+ T cells (*P=*0.0041) compared to their control counterparts. On the other hand, small intestine of antibiotic-treated YBG-recipients showed higher IL10 (*P=*0.0087), IL17 (*P=*0.0159) and IFNγ (*P=*0.041) production, compared to microbiota-depleted controls. Immune cells from large intestine of YBG-treated microbiota-depleted mice produced significantly higher amounts of IL17 (*P=*0.0034) and IFNγ (*P=*0.0039), but not IL10, compared to control counterparts.

### YBG treatment results in increased immune regulatory SCFA production in B6 mice

Since fermentation of non-digestible sugars by colonic bacteria can result in the generation of metabolites including SCFA, fecal SCFA levels in control and YBG treated B6 mice with intact and depleted gut microbiota were determined. As observed in Fig. 5A, significantly higher fecal acetic acid (*P=*0.016), propionic acid (*P=*0.026) and butyric acid (*P=*0.013), and lower valeric acid (*P=*0.036) levels were detected in YBG-treated mice compared to their control counterparts. On the other hand, depletion of gut microbiota not only resulted in the elimination of YBG-treatment associated effect on SCFA production, but caused profound suppression of overall SCFA levels in both control and YBG treated mice (Fig. 5B) compared to that of mice with intact microbiota (Fig. 5A).

**FIGURE 5:**
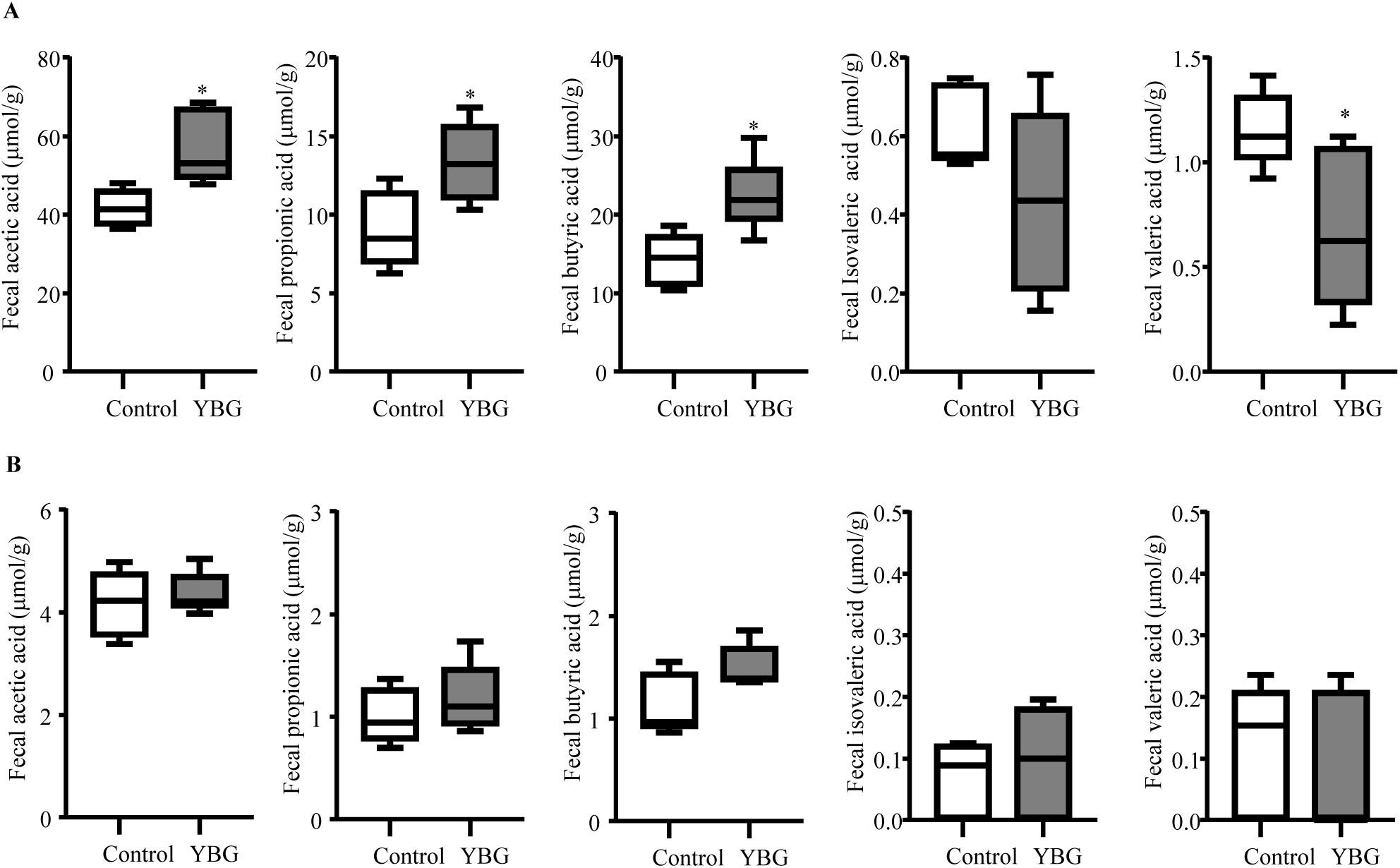
Impact of oral administration of YBG on the fecal SCFA levels in B6 mice with intact (A) and depleted (B) gut microbiota. (A, B) SCFA levels in fecal pellets collected from control and YBG treated mice that were left alone or given broad spectrum antibiotics are shown. Mean±SDs (*n*=5) are shown for all panels. *Different from control, *P*<0.05.

### YBG treatment associated protection of B6 mice from colitis is gut microbe-dependent

To determine if the YBG treatment associated protection of B6 mice from colitis is microbiota dependent, YBG treated and control B6 mice were given broad spectrum antibiotics to deplete gut microbiota and given DSS to induce colitis as depicted in **Fig. 6A**. As observed in **Fig. 6B and C**, control and YBG treated mice with intact gut microbiota showed weight loss and disease severity comparable to that of Fig. 1. Importantly, control mice with microbiota depletion showed relatively lesser weight loss and disease severity compared to their control counterparts, and the difference was statistically significant (*P=*0.012) only at 10 week time point (**Fig. 6B; left panel**). However, colon inflammation scores were not statistically significant in these mice (**Fig. 6C; left panel**). On the other hand, YBG-treated, microbiota-depleted mice showed significantly higher weight loss and disease severity starting at day 6 time-point (*P=*0.016) and more severe thereafter (*P<*0.01) compared to untreated mice with intact microbiota. Further, YBG treated, microbiota depleted mice showed higher colon inflammation (*P=*0.047) compared to control mice (**Fig. 6C; right panel**). Examination of cytokine profiles of immune cells from the colons revealed significantly higher production of pro-inflammatory IFNγ (*P*<0.001) and IL17 (*P*<0.001), and comparable anti-inflammatory IL10, production in YBG-treated, microbiota depleted mice compared to YBG treated mice with intact gut microbiota (**Fig. 6D**). However, cells from microbiota depleted control mice showed diminished IFNγ production compared to cells from controls with intact microbiota (*P*<0.001).

**FIGURE 6:**
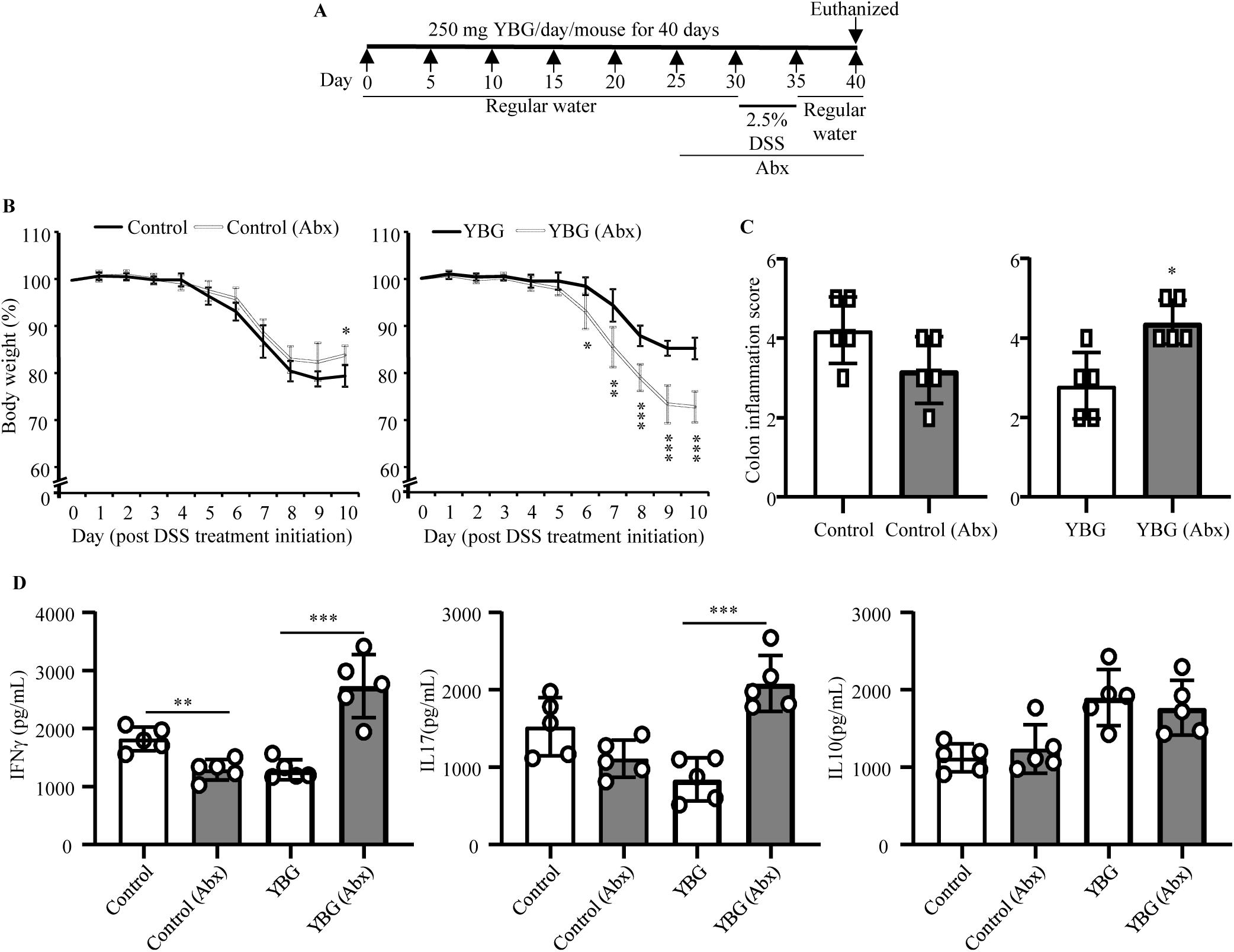
Effect of microbiota depletion (A) in DSS colitis induced weight loss (B), colon inflammation (C) and cytokine secretion by intestinal immune cells (D) in control and YBG treated B6 mice. (A) Cartoon depicting experimental design is shown. (B) Body weights of individual animals were measured every day starting at day 30 and changes in percentage of body weights, relative to initial body weight, are shown. (C) H&E stained distal colon sections were evaluated for inflammation and the severity scores are shown. D) Concentrations of indicated cytokines secreted by colonic immune cells upon *ex vivo* stimulation using anti-CD3 antibody are shown. (B,C, D) Mean±SDs (*n*=5) are shown. *, **,*** Different from control, *P*<0.05, *P*<0.01, *P*<0.001 respectively.

## Discussion

Literature suggests that certain complex dietary polysaccharides such as BGs of different microbial and plant origin affect the host immune function (44–48). While previous studies using various BG preparations and BG rich diets have shown that they have the ability to suppress gut inflammation in colitis models (10–14), others have shown that BGs aggravate the colitis severity (15). Further, while it has been shown that genetic deletion of a BG interacting receptor, Dectin-1, does not affect the course of murine experimental colitis, another group found Dectin-1 deficiency increased susceptibility to colitis (16, 17). Hence, although BGs from different sources can vary structurally and functionally (1), the true impact of orally administering BGs on gut inflammation and the associated mechanisms is unknown, and the previously reported outcomes are non-conclusive without the use of well-defined, high pure, BGs such as the YBG used in this study. Here, we show that prolonged oral administration of high pure YBG prior to DSS treatment to induce colitis results in significantly lowered severity of gut inflammation in B6 mice. This diminished susceptibility to DSS colitis is associated altered gut microbiota composition and function favoring polysaccharide metabolism, higher immune regulatory SCFA production, and enhanced gut immune regulation.

Intriguingly, we found that treatment with high pure YBG at disease induction stage alone has no beneficial effects, but only prolonged oral pretreatment with this agent suppresses colitis susceptibility. This shows that well defined BGs can promote protection from colitis, but it depends on gradually acquired enhanced immune regulation, perhaps mediated by altered / YBG shaped gut microbiota. In fact, although the role of immune regulation promoted directly by YBG in the gut mucosa cannot be ruled out, we found that prolonged oral treatment using YBG causes changes in the gut microbial composition. YBG consumption resulted in reduced abundance of Firmicutes and increased abundance of Bacteroidetes and Verrucomicrobia phyla, which include many polysaccharide fermenting bacteria (49, 50). However, YBG treatment also diminished gut microbial diversity / species richness, suggesting that YBG treatment induced effect is due to selective enrichment of microbial communities that promote immune regulation, but not by increasing the diversity of gut microbiota. Further, predictive functional profiling of fecal microbiota showed overrepresentation of metabolic pathways linked to carbohydrate metabolism, glycan biosynthesis and metabolism, and biosynthesis of secondary metabolites after YBG treatment compared to pre-treatment time-point. More importantly, our novel observations that fecal levels of immune regulatory microbial SCFA (51, 52) such as acetic acid, propionic acid and butyric acid produced by gut microbiota are higher in YBG treated mice suggest that altered microbiota is functionally superior in terms of their ability to promote immune regulation. This notion has been validated by diminishes levels of these SCFAs, Treg frequencies in the gut mucosa, and elimination of YBG treatment induced effect upon depletion of gut microbiota. Further, the need for prolonged pre-treatment, which could potentially help gradual shaping of gut microbiota structure and function, for protection from colitis is another indication that altered gut microbiota is the key contributor of protection from colitis in YBG fed mice.

Colonic microbial communities can ferment non-digestible dietary polysaccharides to use them as energy sources for their growth, and also produce small metabolites such as SCFA that enhance gut immune regulation (16, 49, 51, 53–58). Previous studies have shown that gut colonization by specific commensal microbial communities causes induction and expansion of Foxp3+ Tregs in the intestinal and systemic compartments (59–61). Further, gut bacteria play a critical role in maintaining peripheral tolerance and gut immune homeostasis (62, 63). However, it is not known if BG shaped gut microbes promote Treg induction and/or expansion. Our observations that YBG treatment increases the production of microbial SCFA, which are known to enhance gut integrity and immune regulation (51, 55), suggest that microbiota does contribute to YBG treatment associated increase in overall immune regulation. Importantly, abundances of microbes belonging to polysaccharide degrading and well as host beneficial communities such as *Parabacteroides, Sutterella, Prevotella, Bifidobacterium and Akkermansia* are increased upon prolonged YBG treatment. Moreover, microbial depletion and reduced SCFA production resulted in increased production of pro-inflammatory cytokines IFNγ and IL17 in the distal gut of YBG treated mice substantiating the notion that microbes contributes to enhance immune regulation and diminished susceptibility to colitis when pre-treated with this CDP.

Although this study is focused on the impact of YBG treatment on colitis susceptibility and distal gut, we found that, unlike the large intestine, small intestine of YBG treated mice, with intact and depleted microbiota, have relatively higher frequencies of Tregs compared to untreated controls. This suggests that YBG-treatment associated increase in Treg abundance in small and large intestinal compartments may involve different mechanisms. It is possible that while prolonged treatment with YBG, at the employed human relevant dose, interacts directly with small intestinal mucosa to enhance immune regulation, YBG treatment associated enhanced immune regulation in the large intestine may be primarily microbiota (and SCFA) dependent. Of note, our recent report showed that treatment of mice with a higher dose of this YBG caused strong pro-inflammatory immune response in both small and large intestine (29), suggesting that use of higher doses of this CDP is not advisable to ameliorate susceptibility to gut inflammation.

Although we previously reported that Dectin-1 dependent interaction of high pure YBG with immune cells could promote a combination of both pro-inflammatory and immune regulatory immune responses (19, 27) under normal circumstances, our current study suggest that such direct interaction can produce primarily pathogenic response under depleted microbiota and pro-inflammatory conditions such as a chemical insult using DSS. In fact it has been shown that the degree and type of immune cell response to BGs depends on the cytokine milieu and microenvironment (64). It is possible that YBG is degraded by colonic bacteria and produce immune regulatory SCFA along with minimizing its direct interaction with large intestinal mucosa. On the other hand, microbial depletion could diminish YBG degradation and SCFA production. Further, DSS which causes epithelial damage can expose immune cells for gut mucosa for direct interaction with YBG resulting in dominant pro-inflammation immune response. This is evident from our observation that YBG treated, microbiota depleted mice presents more severe colitis compared to YBG treated mice with intact microbiota, and control groups of mice with intact and depleted microbiota.

Overall, our studies using YBG found that this CDP can promote gut immune regulation by altering gut microbiota, increasing SCFA production, enhancing the overall immune regulation and gut health under normal circumstances suggesting the prebiotic value of BGs. Importantly, only pre-conditioning of the intestinal microenvironment in terms of altering gut microbiota composition and function, which enhances SCFA production and immune regulation, by treating with YBG diminishes susceptibility to chemical induced colitis. We show that YBG treatment has no benefits in ameliorating the severity of ongoing gut inflammation, as in the case of UC. Hence, with respect to the colitis, health benefits of YBG could be limited to those who are susceptible to, but not yet developed, the disease rather than those who have established disease. We employed microbiota depletion approach in our study primarily to understand the mechanism by which YBG treatment enhances gut immune regulation and diminish colitis susceptibility. Nevertheless, our observations suggest that consumption of YBG, as a dietary supplement, by those who are under antibiotic therapy could produce adverse effects such as gut inflammation. Of note, utility of antibiotic treatments have been considered for ameliorating the severity of gut inflammation and for managing UC symptoms (65, 66). Our observations suggest the possibility that consumption of YBG as a dietary supplement by UC patients who are under such therapy will result in aggravated, rather than ameliorated, disease severity. In conclusion, this study, which used high pure YBG, not only demonstrates the dietary supplement value of BGs in promoting gut health and the associated mechanisms, but also show that these CDPs may not be beneficial in suppressing ongoing gut inflammation and they produce adverse effects when used in combination with antibiotic therapy.

## Conflict of Interest statement

Authors do not have any conflict(s) of interest to disclose.

## Author contribution

R.B., B.J, and J.S. researched and analyzed data, and R.G. researched and analyzed the data and edited the manuscript; C.V. designed experiments, researched and analyzed data, and wrote/reviewed/edited manuscript.

## Acknowledgments

This work was supported by internal funds from MUSC, National Institutes of Health (NIH) grants: R21AI133798 (NIAID), and R21AI133798 and R21AI136339-02S1 Administrate supplements (Office of Dietary Supplements) to CV. The authors are thankful to the Histology core of Pathology department of MUSC for the histology service, and Genomic center of MUSC for 16S rRNA gene sequencing service. Dr. Vasu is the guarantor of this work and as such, has full access to all the data in the study and takes responsibility for integrity of the data and accuracy of data analysis.

## Notes

**Conflict of Interest**: Authors do not have any conflict(s) of interest to disclose.

#### Summary of Updates

Re-submission version of the manuscript

