## Supplemental data for "Pretreatment with yeast derived complex dietary-polysaccharide leads to suppressed gut inflammation, altered microbiota composition and increased immune regulatory short-chain fatty acid production in C57BL/6 mice"

Supplemental Fig. 1

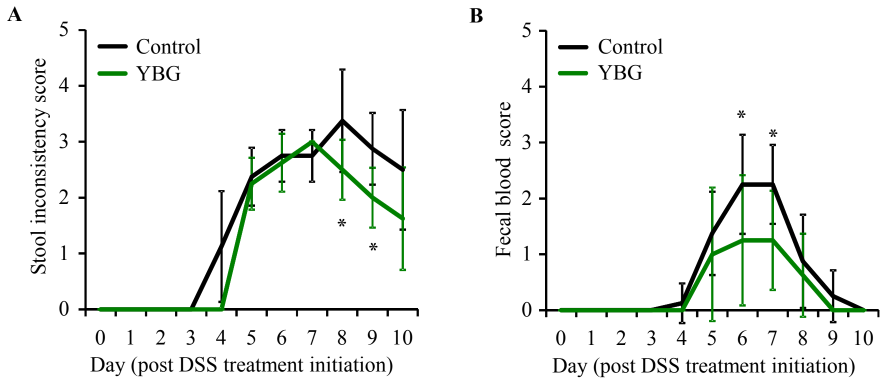


Supplemental Fig 1: Impact of pre-treatment with YBG on DSS induced colitis associated stool softening (A) and fecal blood levels (B). Stool consistency and blood content in stool of mice described in Fig. 1B were monitored daily starting day 30 and scored. Mean±SDs (*n*=8) are shown for all panels. *Different from control, P<0.05.

Supplemental Fig. 2

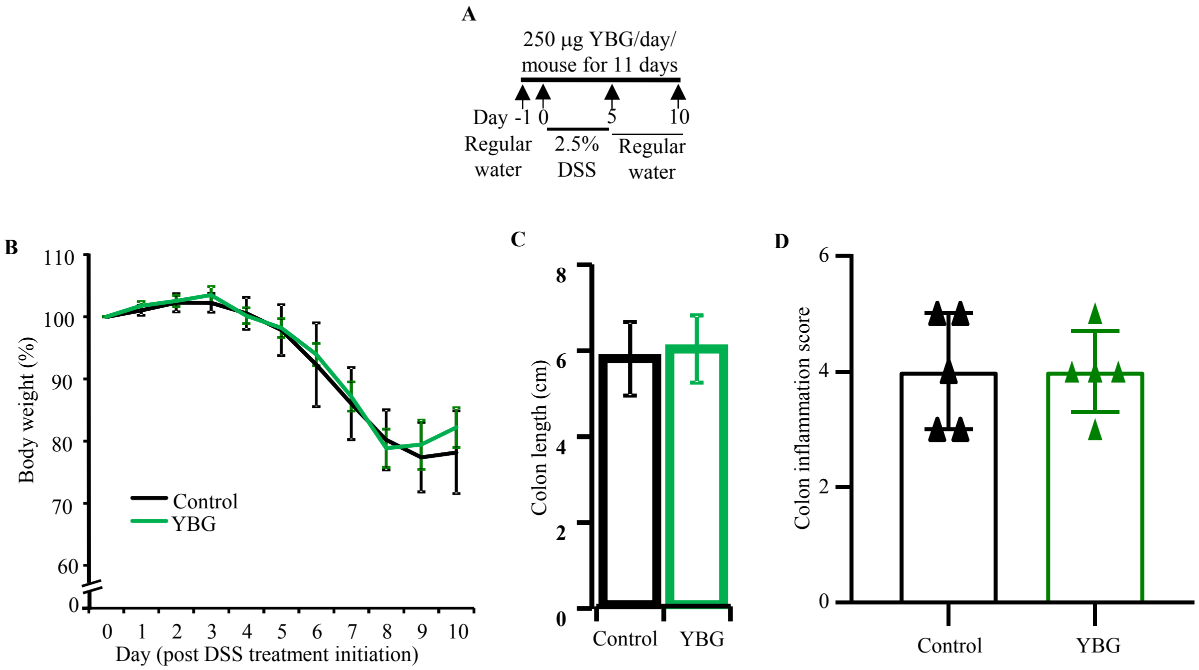


Supplemental Fig 2: Impact of YBG treatment initiation during DSS treatment (A) on colitis associated weight loss (B), shortening of colon (C) and colon inflammation (D) in B6 mice. (A) Cartoon depicting experimental design is shown. B) Body weights of individual animals were measured every day and changes in percentage of body weights, relative to initial body weight, are shown. (C) colon length (right panel) of euthanized mice are shown. (D) H&E stained distal colon sections were evaluated for inflammation and severity scores are shown. (B, right panels of C, D) Mean±SDs (*n*=5).

Supplemental Fig. 3

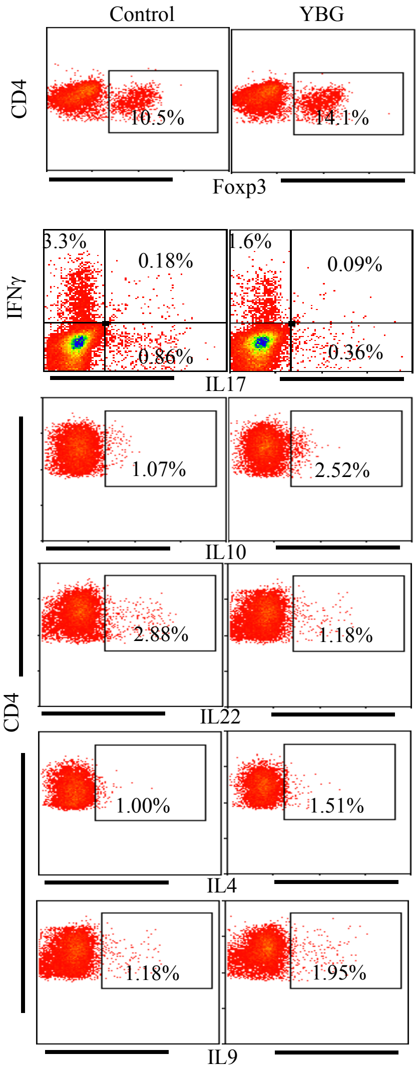


Supplemental Fig 3: Representative FACS plots showing frequencies of cells positive for indicated specific factors. Mean±SD values are shown in Fig. 2; panels A and B.

Supplemental Fig. 4

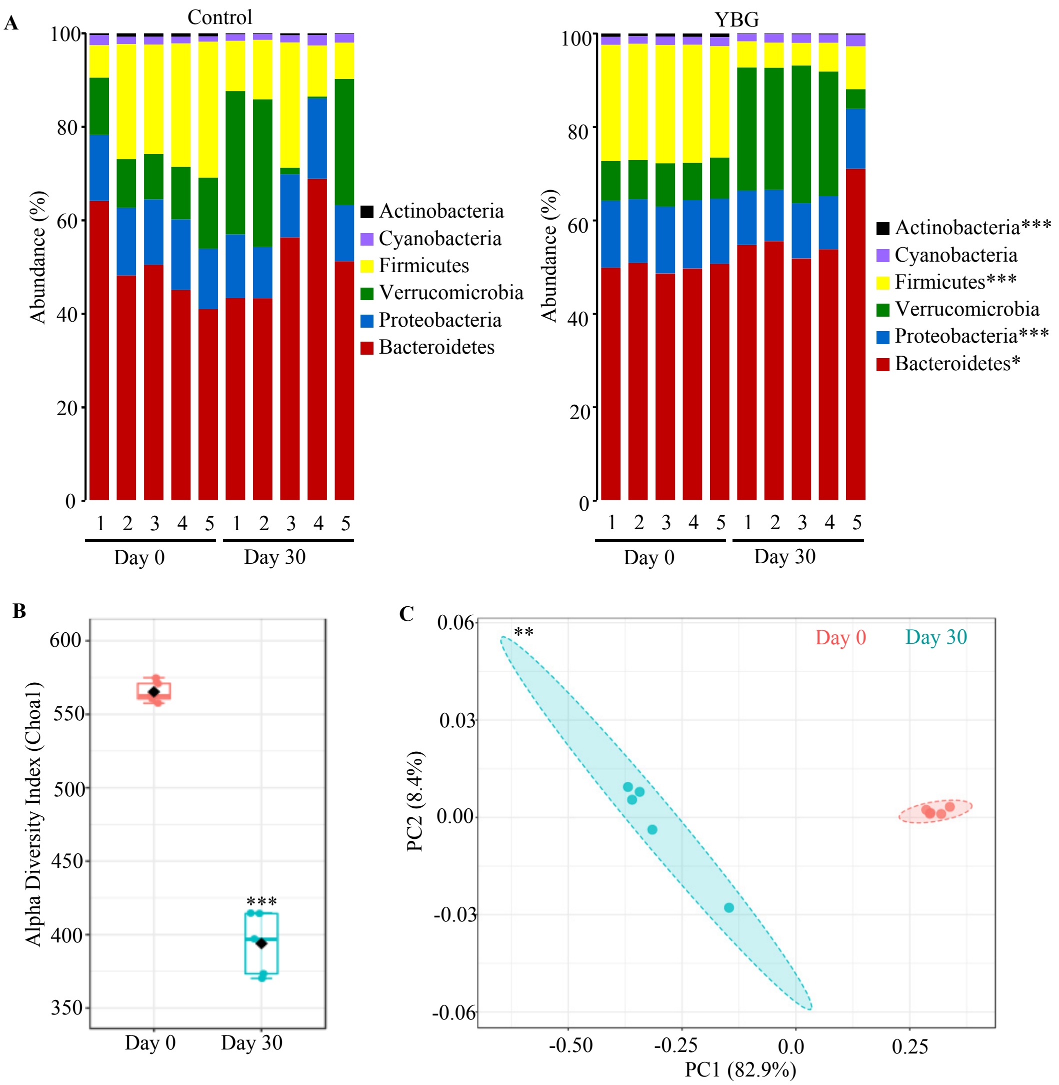


Supplemental Fig. 4: Impact of oral administration of YBG on fecal microbiota composition at phylum level (A), α-diversity/species richness (B) and β-diversity (C) in B6 mice. (A) OTU data of fecal samples collected from control (left panel) and YBG treated (right panel) mice at days 0 and 30 described for Fig. 3 were compiled to phylum level. (B) α-diversity/species richness of 16S rRNA gene sequences of samples collected on days 0 and 30 from YBG treated mice. (C) Principal component (PC)/β-diversity measures of fecal bacterial communities of YBG treated mice. n=5 for all panels. *,**,*** Different from control; *P*<0.01, P<0.001 respectively.

Supplemental fig. 5

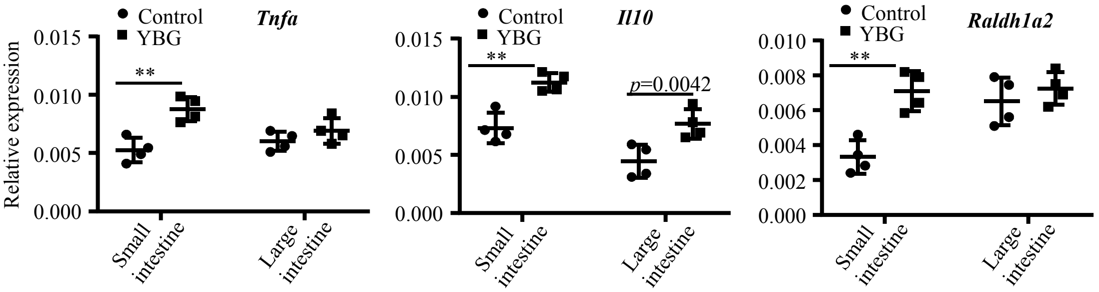


Supplemental Fig. 5: Effect of oral administration of YBG on cytokine expression in the intestine of B6 mice. cDNA prepared from the distal ileum and distal colons of YBG treated and control mice were subjected to qPCR and the expression levels of cytokines and non-cytokine factors relative to β-actin expression levels were compared. Mean±SDs ($n$=4) are shown. ** Different from control; $P$<0.01 respectively.

Supplemental Fig. 6

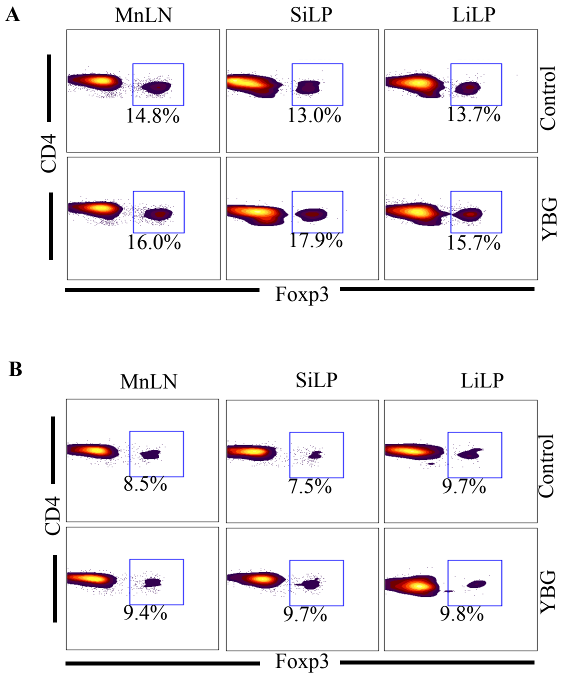


Supplemental Fig 6: Representative FACS plots showing frequencies of Foxp3+ cells in control and YBD treated B6 mice with intact (A) and depleted (B) gut microbiota. Mean±SD values are shown in Fig. 4 A and B, left panels.

Supplemental Fig. 7

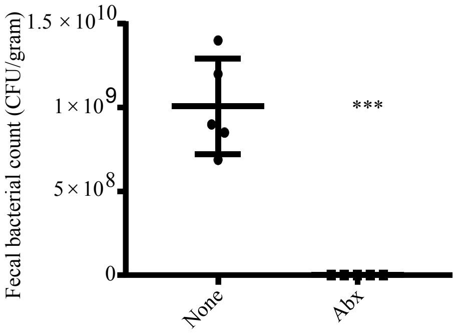


Supplemental Fig. 7: Impact of treatment with antibiotic cocktail (Abx) on fecal microbes in B6 mice. Fecal pellets collected from individual mice were suspended in sterile PBS, plated onto brain heart infusion plates under anaerobic and aerobic conditions for up to 72 h and the total number of colonies (colony forming units; CFU) were determined. Mean±SDs ($n$=5) are shown. *** Different from control; $P$<0.001 respectively.
